# Partial RNA Design

**DOI:** 10.1101/2023.12.29.573656

**Authors:** Frederic Runge, Jörg K.H. Franke, Daniel Fertmann, Rolf Backofen, Frank Hutter

## Abstract

RNA design is a key technique to achieve new functionality in fields like synthetic biology or biotechnology. Computational tools could help to find such RNA sequences but they are often limited in their formulation of the search space. In this work, we propose *partial RNA design*, a novel RNA design paradigm that addresses the limitations of current RNA design formulations. Partial RNA design describes the problem of designing RNAs from arbitrary RNA sequences and structure motifs with multiple design goals. By separating the design space from the objectives, our formulation enables the design of RNAs with variable lengths and desired properties, while still allowing precise control over sequence and structure constraints at individual positions. Based on this formulation, we introduce a new algorithm, libLEARNA, capable of efficiently solving different constraint RNA design tasks. A comprehensive analysis of various problems, including a realistic riboswitch design task, reveals the outstanding performance of libLEARNA and its robustness.

## 1 Introduction

RNA design describes the problem of generating RNA sequences with certain properties to fulfill desired functions [Hammer et al., 2019]. The design endeavors often start from carefully selected RNA fragments with known functions to combine these so-called motifs and achieve new functionality [Wachsmuth et al., 2012, Li et al., 2018, Nozawa et al., 2019]. The designed construct is then evaluated in a wet laboratory experiment proofing the desired functionality. To save costs, however, computational methods are employed to reduce the number of initial candidates for experimental validation [Hammer et al., 2019].

The theoretical design space for computational models is typically huge. Thus, restricting this space to a reasonable subspace can drastically speed up the search for good candidates. It is common knowledge that primary- and secondary structure features are useful restrictions, while the design goals might vary depending on the particular application [Hammer et al., 2019]. To optimally support RNA practitioners, an algorithm should thus be able to process any primary- and secondary structure restrictions, efficiently explore large design spaces, consider different design goals, and provide as many candidates as needed, optimally with different lengths and compositions. We observe that current RNA design approaches do not fully satisfy these requirements. For example, consider the design of constructs that contain a GNRA tetra-loop motif to establish tertiary interactions for subsequent crystallization studies [Reyes et al., 2009]. Then one might want to design 50 candidate constructs with variable lengths, that contain a stem of unspecified length with a GNRA tetra-loop. This kind of task cannot be tackled with any current method, because they either cannot design sequences with different lengths [Hofacker et al., 1994, Kleinkauf et al., 2015, Runge et al., 2019, Minuesa et al., 2021], cannot consider unconstrained regions [Andronescu et al., 2004, Reuter and Mathews, 2010], or cannot handle designs with unknown shapes [Avihoo et al., 2011, Drory Retwitzer et al., 2016]. We are not even aware of any formulation that supports the definition of a task with a single paired position without defining its pairing partner, region, or an entire final shape because design approaches do not consider unbalanced pairing schemes.

In this work, we propose *partial RNA design*, a novel RNA design paradigm that tackles these issues. Our formulation separates the definition of the design space from the design objectives, allowing arbitrary sequence and structure restrictions of the design space. Using this problem definition, we enhance a previous RNA design approach [Runge et al., 2019] to develop *libLEARNA*, a new algorithm that can explore large RNA design spaces guided by sequence and secondary structure motifs to yield large amounts of variable-length candidate constructs for different design objectives. Our main contributions are as follows:

- We propose *partial RNA design*, a novel RNA design paradigm formulated as a constraint satisfaction problem. Partial RNA design is more general than any previous RNA design formulation and allows the design of RNA sequences with different lengths while providing precise control over local and global restrictions of the sequence and structure space (Section 2).
- We improve an existing RNA design framework, LEARNA [Runge et al., 2019], with a masked training objective to develop *libLEARNA*, a new algorithm that is capable of efficiently navigating large RNA design spaces, ensuring the inclusion of desired motifs while exploring the space within the set parameters (Section 3).
- We evaluate *libLEARNA* on various RNA design tasks, demonstrating its strong performance and robustness across multiple search spaces and design objectives (Section 4).
- We open-source our RNA design framework at https://github.com/automl/learna_tools.

## 2 Partial RNA Design

In this section, we formulate partial RNA design as a constraint satisfaction problem (CSP). CSPs are general formulations of combinatorial problems defined as a triple (*X, D, C*), where *X* is a set of variables, *D* is a set of their corresponding domains of values, and *C* is a set of constraints. Each variable can take on the values of its corresponding non-empty domain. A constraint *c ∈ C* is a tuple (*t, R*) with *t ⊆ X* being a subset of *k* variables from *X* and *R* being a *k*-ary relation over these variables. An *evaluation* of the variables is a function from a subset of variables to a set of values from their corresponding domains that satisfies a constraint (*t, R*) if the assigned values to the variables in *t* satisfy the relation *R*. An evaluation is called *consistent* if it does not violate any of the constraints and *complete* if it includes all variables in *X*. A solution to the respective CSP is an evaluation that is consistent and complete.

While CSP formulations for modeling RNA design exist [Garcia-Martin et al., 2013, 2015, Minuesa et al., 2021], the variables of these CSPs correspond to individual positions in an RNA. In contrast, partial RNA design considers an RNA design space that can be restricted at certain regions. A region consists of a primary- and secondary structure part, each represented with an individual variable in our CSP formulation. We dub the tuple of two corresponding primary- and secondary structure parts, i.e., parts that describe the same region of a design space, a *motif*. We allow defining unconstrained positions within each part of a motif by introducing two wildcards: A single position wildcard (denoted ? in our CSP formulation) that defines an unconstrained position in a part of a motif, and a variable-length wildcard (denoted 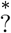 in our formulation) that marks a position that can be arbitrarily extended, i.e., we can insert any sequence of primary- or secondary structure symbols at the respective position. This allows us to define arbitrarily restricted design spaces of variable length. For example, recall our example of the design of sequences that contain a GNRA tetra-loop. Using the common dot-bracket notation [Hofacker et al., 1994] we can define the design space as follows. We define the sequence part of the loop region as `?GNRA?`, with a corresponding structure part (….) using two single position wildcards. By adding a variable-length wildcard at both ends, we result in the following design space formulation:

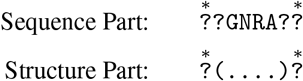

The variable-length wildcards allow to place any sequence of nucleotide or structure symbols at the respective position, while the matching pair of brackets ensures the formation of a stem with a GNRA loop. Desired objectives can be added independently on top of the design space formulation via constraints, which separates the formulation of the design space from the objectives. For partial RNA design, we use the common dot-bracket notation to denote secondary structure features but note that a generalization of partial RNA design to different alphabets and molecule classes is rather straightforward.

### 2.1 The Partial RNA Design Problem

To generalize RNA design to partial RNA design, we first extend the RNA nucleotide and secondary structure alphabets, Φ := { A, C, G, U } and Ω := { .,), (, }, with a *wildcard symbol* “?” indicating a position that is not constrained. Similarly, we add a *variable length wildcard symbol* 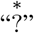 to indicate an unconstrained region of variable length:

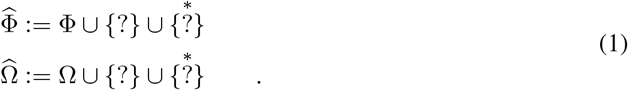

*We define a motif* **m** as a pair 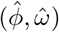, consisting of an RNA sequence part 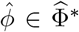, and an RNA structure part 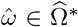, where *\** denotes the *Kleene closure* [Fletcher et al., 1991]. *At each position* 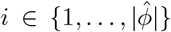 of the sequence part, 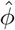, we can define a set of valid assignments, 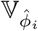, for the particular position *i*:

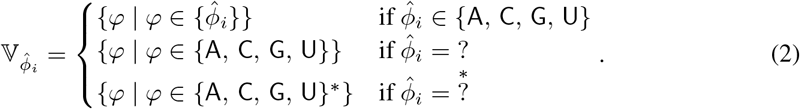

Note that *Φ* can denote a single symbol or a sequence of symbols, depending on whether the respective position *i* of 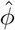 is assigned a variable length wildcard or not. A valid assignment *ϕ* ∈ Φ*\** for the entire sequence part of the motif is then the concatenation (○) of a single element picked from each set of valid assignments at each position,

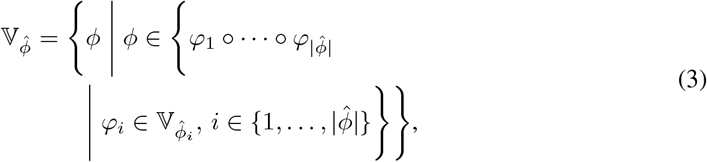

Similarly, for the structure part, 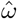, of the motif **m**, we define the sets of valid assignments 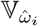 and 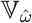 as

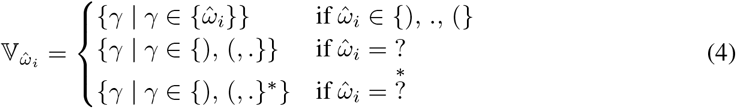

and

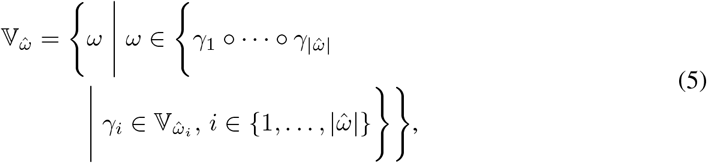

with 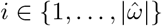. For the entire motif **m**, the set of valid assignments, 𝕍**m**, can then be written We define as follows:

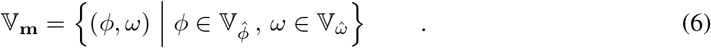

two variables, *x* and *y*, for the sequence and the structure part of a motif, respectively. The corresponding domains of values for *x* and *y* are defined as

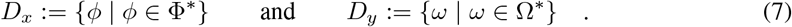

To meet the constraints given by the motif **m**, the assigned values for *x* and *y* have to be in the set of valid assignments. We define the relation 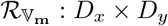 as

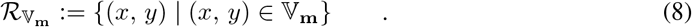

For a sequence, 𝕊= (**m**1, …, **m***k*), *k* ∈ ℕ, of desired motifs, we use two variables *xi* and *yi, i* 1, …, *k*, for each motif. These variables will define the set of variables *X* of our CSP formulation for partial RNA design. We can think of the motifs in 𝕊 as follows: When we pick a valid assignment for each motif from the sets of valid assignments, the concatenation of the sequence parts defines a designed RNA primary structure, while the concatenation of the structure parts defines a dot-bracket annotated secondary structure. All possible combinations of single elements picked from all sets of valid assignments thus define the entire design space of the motifs in 𝕊. However, the secondary structures are not necessarily valid structures, since we do not restrict the assignments in any way and the variables are independent of each other. We address this with the definition of objectives. An objective *O*((*x*1, …, *x*_*k*_, *y*_1_, …, *x*_*k*_)) defines a relation between the variables *xi* and *y*_*i*_. The solution space of a given objective *O* defines the set of tuples 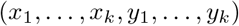 that satisfy the objective. We can write this as the relation 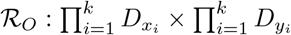, with

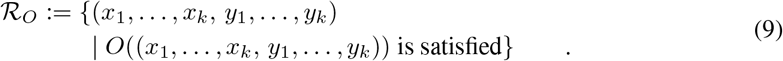

For example, to define a global folding relation, we can define the relation *ℛ*_ℱ_ over the concatenated sequence parts, *x*_1_ ○ · · · ○ *x*_*k*_, that have to fold into the concatenation of the structure parts: *ℱ* (*x*_1_ *○ · · · ○ x*_*k*_) = *y*_1_ *○ · · · ○ y*_*k*_. We formulate the relation 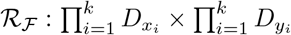, with

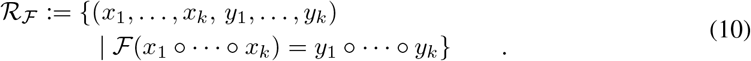

The *partial RNA design problem* can then be defined as follows.

#### Definition 1

***Partial RNA Design Problem*** *Given a sequence of RNA motifs, 𝕊*= (**m**_1_, …, **m**_*k*_), *of length k ∈ ℕ and a set of n ∈ ℕ objectives 𝕆* = {*O*_1_, …, *O*_*n*_}, *a solution to the partial RNA design problem associated with the sequence 𝕊 corresponds to a solution of the constraint satisfaction problem* (*X, D, C*), *defined as follows:*

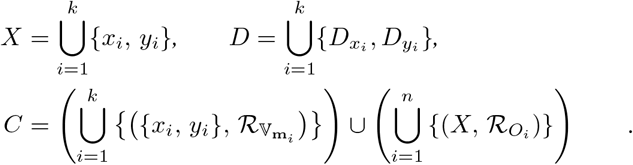

The partial RNA design problem describes a completely novel RNA design paradigm: In contrast to previous work that tries to find optimal solutions in a very restricted setting, our approach designs RNAs that are contained in a broader design space, essentially designing RNA focus libraries while considering arbitrary primary- and secondary structure constraints. Furthermore, partial RNA design covers multiple existing problem formulations: Inverse RNA folding can e.g. be defined with a single motif of a complete secondary structure with an unknown sequence. We also note that we do not limit the definition of the motifs. For example, structure parts can contain any composition of brackets, balanced or unbalanced. This allows us to create structural diversity and further enables us to extend the task at any given point since we are no longer bound to explicit pairing positions. However, one can define an objective to mark specific positions that have to be paired. In the following section, we detail the development of libLEARNA, an algorithm that is capable of designing RNAs from provided sequence and structure motifs under different objectives.

## 3 Method

In this section, we develop libLEARNA, a new algorithm capable of solving the partial RNA design problem. Partial RNA design can be interpreted as a masked prediction task, where the goal is to assign values to the unknown regions of a partially restricted RNA design space, similar to the masked training objective of e.g. the language model BERT [Devlin et al., 2018]. For libLEARNA, we seek to learn the navigation of a partially restricted design space across thousands of tasks with fixed lengths. During evaluation, a task of variable length, i.e., a task that contains a variable length wildcard symbol 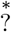, is interpreted as a set of fixed-length tasks. To solve it, we sample a fixed-length task from the provided design space and try to solve it with a single shot. Since partial RNA design separates the objectives from the formulation of the design space, this strategy imposes three challenges: (i) Our algorithm has to be capable of navigating partially restricted RNA design tasks, it has to be able to adapt to changing objectives, and (iii) it requires strong performance with the first prediction. Before we dive into the details of how to achieve these goals, we briefly recap the foundation of libLEARNA, LEARNA [Runge et al., 2019].

LEARNA is an automated deep reinforcement learning (AutoRL) [Parker-Holder et al., 2022] algorithm for the generative design of RNA sequences, following an inverse RNA folding approach [Hofacker et al., 1994]. The algorithm has two stages: offline training & optimization and online application. At offline training time, in an inner-loop, an agent learns an RNA design policy via periodical interactions with an environment. The agent receives a state from the environment and chooses an action based on the current state. The action is then communicated to the environment to compute a reward signal that informs the agent about the value of the chosen action. In LEARNA, states are defined as local representations of the input structure and actions correspond to placing a nucleotide or placing Watson-Crick pairs in case the structure indicates that the given position is paired. The reward is computed once all sites have been assigned nucleotides; after applying a folding algorithm to the designed sequence, the reward function is based on the Hamming distance between the folded candidate and the desired structure. The policy of the agent is defined through a neural network.

After training, still as part of the offline phase, the agent is evaluated on a validation set, and the loss is communicated to an efficient Bayesian Optimization method, BOHB [Falkner et al., 2018], that jointly optimizes the configuration of the RL system in the outer loop, seeking to minimize the observed loss. During online application to a new test problem, the configuration that worked best on the validation set is then executed. Runge et al. [2019] propose three versions within this framework: (1) A version that is not trained at all but updates its policy during evaluation to adapt to a task at hand (LEARNA), (2) a version that leverages the pretraining of the policy but fixes the weights during evaluation for fast sampling of candidates (Meta-LEARNA), and (3) a pre-trained version that allows weight updates at evaluation time (Meta-LEARNA-Adapt). We use the Meta-LEARNA-Adapt approach to develop libLEARNA, because it leverages the advantages of a pre-trained policy while being able to adapt to a given task at hand.

### 3.1 A Decision Process for Partial RNA Design

A reinforcement learning problem can be defined as a Markov Decision Process *𝒟* := (*𝒮, 𝒜, ℛ, 𝒫*), with a set of *states 𝒮*, a set of *actions 𝒜*, a *reward function ℛ*, and a *transition function 𝒫*. For our formulation, we consider a sequence of fixed-length motifs 𝕊 = (**m**1, …, **m***n*), *n ∈ ℕ* with each motif 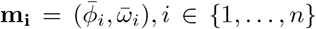, consisting of a sequence part 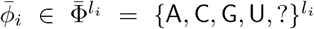 and a structure part 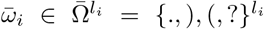 of the same length *li*. Note that, in contrast to 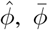 contains no variable length wildcard 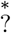. For simplicity, we define the task *τ* as the concatenation *○* of the sequence and structure parts of the motifs: 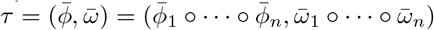.

At each time step *t* = 0, 1, 2, …, *T*, where *T* denotes the terminal time step of the episodic interaction between the agent and the environment, the environment provides a state *s*^*t*^ ∈ 𝒮. The agent chooses an action *a*^*t*^ ∈ 𝒜 based on the provided state and the environment transitions into a new state *s*^*t*^ +1 based on the chosen action, following the transition dynamics 𝒫. At the final time step *T*, the environment computes a scalar reward defined by the reward function ℛ that is communicated to the agent to guide the learning of a policy. For libLEARNA, the states correspond to local representations of the provided task and actions correspond to placing nucleotides. To provide states, the environment transitions over local representations at unconstrained sites of the sequence part of the task. The length *T* of the interaction between the environment and the agent thus depends on the provided motifs in 𝕊. The reward during training is based on the number of satisfied constraints of the structure part of the task. In the following, we detail the individual components of the decision process *𝒟*.

#### 3.1.1 State Space

While Runge et al. [2019] only considers structure features for the state, we consider primary and secondary structure features for libLEARNA. To enable libLEARNA to process sequence and structure information, we numerically encode pairs of a sequence symbol and its corresponding structure symbol for each position of a given design task by indexing all pairs in the set 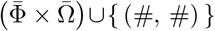. We provide local information about the task to the agent by setting the state *st* to the (2 *κ* + 1)-gram centered around the position of the numerical vector representation of the task *τ*. Since we only consider unconstrained regions, *t* corresponds to the t-th unconstrained position (denoted with ?) of the sequence part 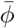. κ is a hyperparameter dubbed the *state radius*. To be able to construct the centered n-gram at all sites, *κ* padding characters (“#”) are introduced at the start and the end of 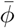 and 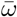. Formally, the state space can be written as

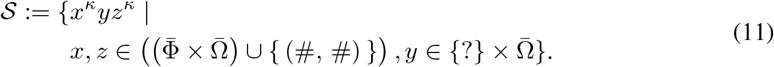

Besides the hyperparameter *κ*, we define a parameter *σ* that decides whether previous actions contribute to the current state or not: If *σ* is set to true, at each time step *t >* 0, the last placed nucleotide replaces the respective wildcard in the nucleotide constraints to inform the agent about its actions. The numerical representation of the task is then updated at each step accordingly.

#### 3.1.2 Actions Space

In each episode, the agent designs an RNA sequence 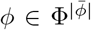, that satisfies all constraints of the sequence part 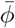. To design a candidate solution the agent places nucleotides by choosing an action *at* at each time step *t* to fill the sites that are unrestricted in 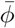. Since an RNA sequence consists of four nucleotides (A, C, G, and U), the action space is formulated as 𝒜:= {0, 1, 2, 3} ≡ {A, G, C, U}. However, we define the *action semantics* hyperparameter that, if active, leverages knowledge about paired sites by directly placing Watson-Crick base pairs (AU, UA, GC or CG) as proposed by Runge et al. [2019]. Note that we do not enforce balanced brackets. For an opening bracket, the site of the corresponding closing bracket thus might not be known. In this case, a single nucleotide is placed for an opening bracket regardless of the choice for the *action semantics* parameter.

#### 3.1.3 Reward Function

The reward function implements the set of objectives 𝕆 of our formulation of partial RNA design (see Definition 1). To be able to navigate large RNA spaces, we need to learn a reasonable relation between the sequence space and the structure space. The folding relation from Equation 10 appears as an objective that relates both in a meaningful way. Therefore, we focus on learning to generate sequences that fold into a structure that satisfies all structure constraints during training and leverage this knowledge at evaluation time when designing RNAs for different objectives. We use RNAFold [Lorenz et al., 2011] with minimum free energy (MFE) structure predictions for folding of the candidate sequences during training.

At the terminal time step *T*, the agent has assigned nucleotides to all sites, and the environment computes the reward ℛ^*T*^. The reward is computed as the number of structure constraints that are satisfied, thus, unconstrained sites are ignored. Formally, we define the structure-loss 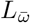 as

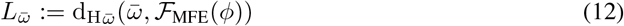

with

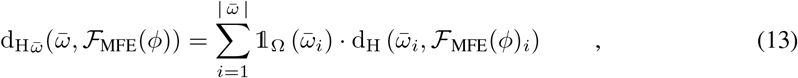

where 𝟙Ω is the indicator function that returns 0 if the i-th position of 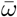 is unrestricted, 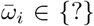, and 1 if 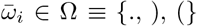, and dH(*·, ·*) is the Hamming distance. We normalize this loss by the length of 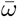 to formulate the reward *ℛ*^*T*^ :

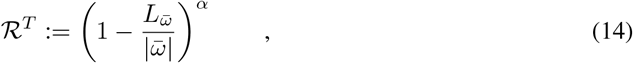

with *α* being a hyperparameter for shaping the reward. We note again that the reward function depends on the objective and we change the reward function at evaluation time accordingly (see Section 4).

#### 3.1.4 Transition Dynamics

At each time step *t*, the state is set to a fixed (2 κ + 1)-gram centered around the position in the numerical representation of the task that corresponds to t-th unconstrained position of the sequence part 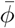. Subsequent states are defined by transitioning to the next unconstrained position. Depending on the choices of the action semantics and the state composition, the transition dynamics may vary and are implemented accordingly.

### 3.2 Training Regimen

We generate three training datasets from all sequences of the Rfam [Griffiths-Jones et al., 2003] database version 14.1 with different length distributions (≤ 200 nucleotides (nt), ≥ 200 nt, random length) of 100000 samples each, and a non-overlapping validation set of 100 samples using *RNAFold* [Lorenz et al., 2011] to receive the structures. We describe all datasets in Table 5 in Appendix A. The choice of the training dataset is a hyperparameter. We use a masked training objective similar to the masked language model training in BERT [Devlin et al., 2018]. The task of *libLEARNA* during training is to fill the masked parts of the sequence such that, after folding the designed sequence, all positions of the folding satisfy the positional constraints of the masked structure. To derive partially restricted tasks, we first mask up to five parts of the structures, each covering up to 20% of the total length, while the positions, the lengths, and the number of parts are sampled uniformly at random. In the second step, we mask corresponding parts of the sequences that remain unmasked in the structures to derive tasks of alternating sequences and structure constraints. Finally, we randomly mask the sequences of ∼ 20% of the samples to derive tasks that correspond to RNA design from arbitrary sequence and structure motifs. Examples of the resulting training tasks are shown in Table 1.

**Table 1:**
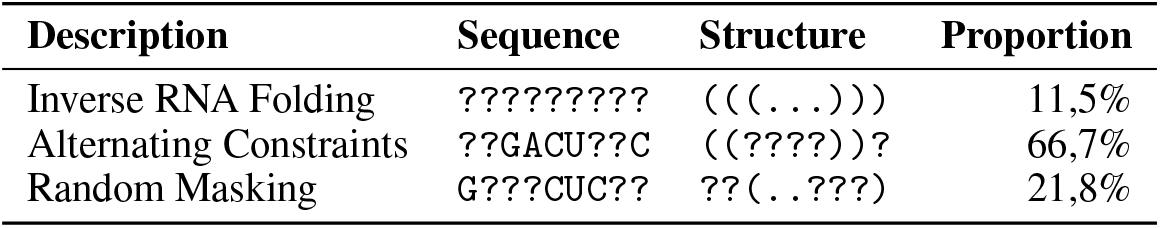
Examples of different tasks in the training data.

### 3.3 Automated Reinforcement Learning

Hyperparameters in reinforcement learning (RL) are known to be sensitive [Henderson et al., 2018]. We, therefore, use an automated reinforcement learning (AutoRL) approach similar to Runge et al. [2019] to automatically find the best RL system for our task in a single algorithm run. Figure 1 outlines the general optimization procedure.

**Figure 1.**
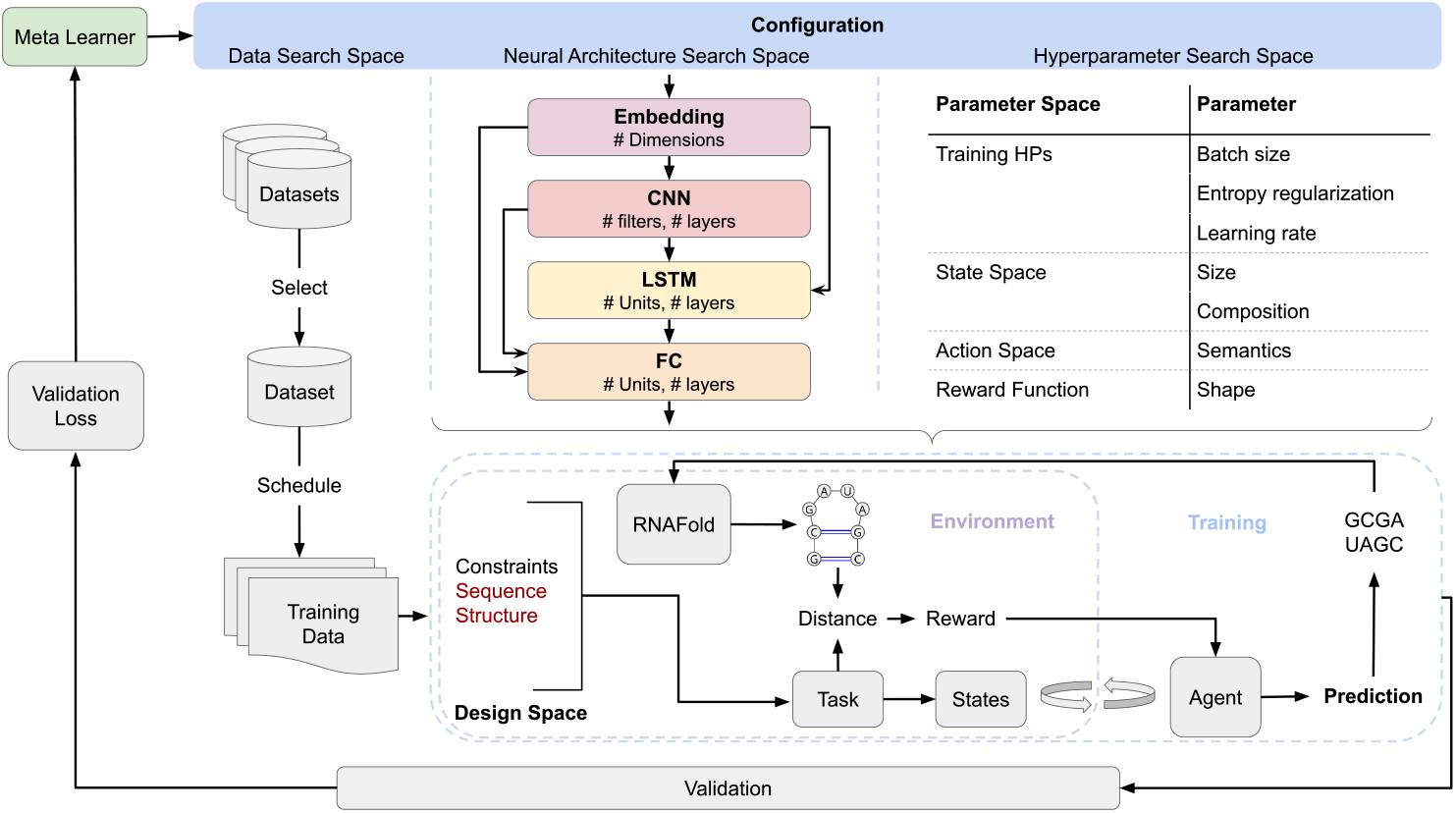
Meta-optimization loop. In each iteration, the meta-learner (BOHB) samples a configuration from a rich configuration space. The sampled configuration defines the learning algorithm’s hyperparameters, a specific environment, a particular network architecture of the agent, a training curriculum, and a training data set. These components together formulate the learner; a deep reinforcement learning algorithm, which is trained on the selected training set and evaluated on a validation set. The resulting validation loss is communicated to the meta-learner to update its model, seeking to learn to sample better configurations with each iteration.

We mainly adopt the configuration space proposed by Runge et al. [2019] but introduce four new dimensions: The *action semantics* parameter to decide whether to predict single nucleotides only or Watson-Crick pairs at paired position, the *individual state composition* parameter to decide if the agent’s actions contribute to the state, as well as two choices that decide about the training data and its schedule. We further allow searching over an additional LSTM layer. The result is an 18-dimensional search space to jointly optimize over the network architecture, all parts of the decision process, as well as training hyperparameters, task distributions, and their schedule, using BOHB [Falkner et al., 2018]. The configuration space, hyperparameters, and the priors we use over them, as well as the final configuration of libLEARNA are shown in Table 6 in Appendix B. We use the exact same setup during meta-optimization as Runge et al. [2019], with the same training budgets and validation protocols. However, while Runge et al. [2019] optimize for an RL algorithm without any policy updates at test time, we directly optimize for an algorithm with policy updates at evaluation time to increase the adaptation capabilities of our approach.

## 4 Experiments

In this section, we show that our proposed strategy is reasonable by demonstrating libLEARNA’s strong performance with the first prediction, its ability to navigate partially restricted design spaces, and its capabilities to adapt to different objectives on tasks with a fixed length (Section 4.1). We then apply libLEARNA to different partial RNA design tasks that require navigation of large design spaces under different objectives (Section 4.2). We use the same model of libLEARNA, obtained from a single meta-optimization run, for all experiments without retraining or changing any parameters.

### 4.1 Validation of Strategy

#### 4.1.1 Improved One-Shot Performance

To analyze the quality of the first predictions of libLEARNA, we compare it to its predecessor with the strongest performance, Meta-LEARNA [Runge et al., 2019], on the 100 inverse RNA folding tasks of the Eterna100 benchmark version 2 [Koodli et al., 2021]. The results are shown in Figure 2.

**Figure 2.**
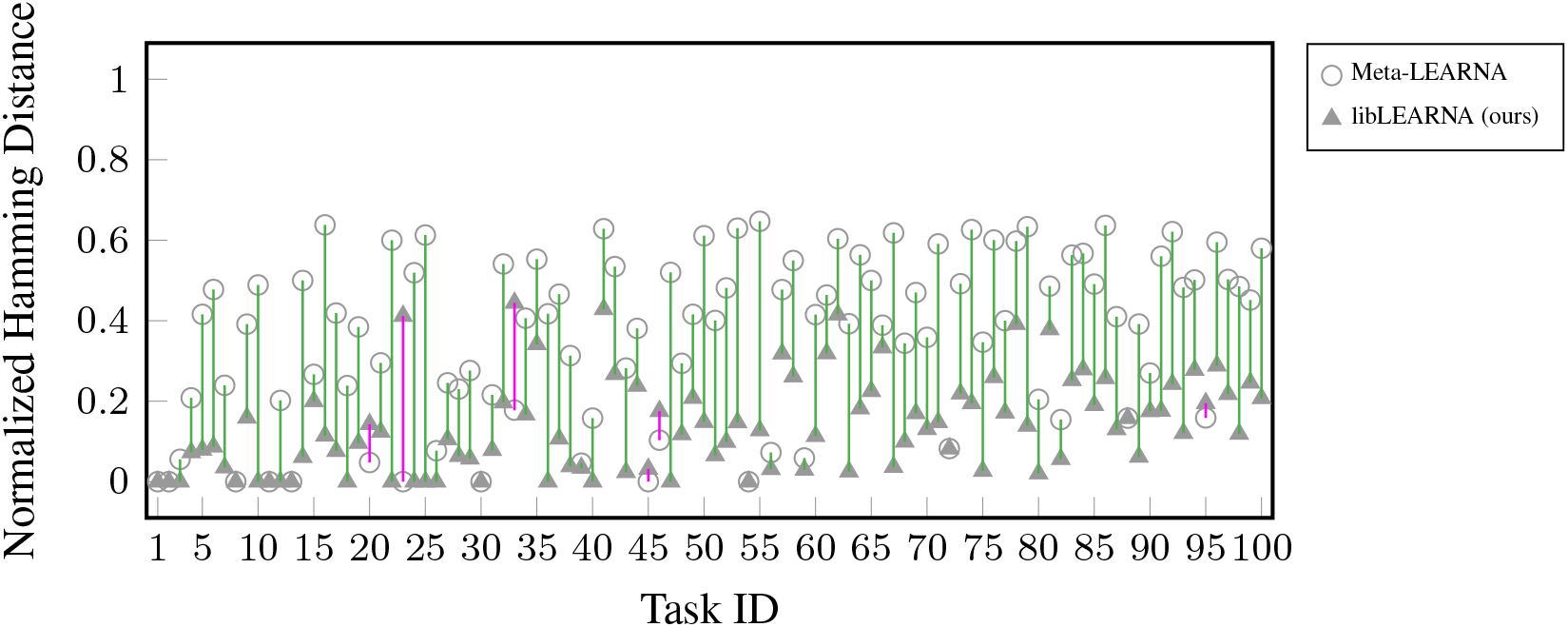
Comparison of the first predictions of Meta-LEARNA and libLEARNA on all tasks of the Eterna100 benchmark version 2. The plot shows the average normalized Hamming distance of the first prediction for all tasks of the benchmark. Green bars indicate tasks where libLEARNA achieves lower Hamming distance, and purple bars indicate tasks where the Hamming distance is above that of Meta-LEARNA’s prediction.

We observe that libLEARNA clearly outperforms Meta-LEARNA, showing a lower Hamming distance for ∼ 95% of the tasks. We take this as an indicator that libLEARNA is generally capable of solving tasks with the first prediction.

#### 4.1.2 Navigating Partially Restricted Design Spaces

To assess libLEARNA’s performance on fixed-length partially restricted tasks, we create a new dataset from the Rfam database version 14.1 and compare libLEARNA against antaRNA [Kleinkauf et al., 2015] and MoiRNAiFold [Minuesa et al., 2021], two algorithms that are capable of predicting RNAs from masked structures of fixed length. We select a total of 100 sequences, ensuring that there is no overlap with the training nor validation data, and randomly mask parts of the sequences and the structures after folding with ViennaRNA’s RNAFold [Lorenz et al., 2011]. In contrast to libLEARNA, antaRNA, and MoiRNAiFold require that all unmasked brackets in the structures are balanced to represent pairs. We therefore unmask the respective brackets if only one part of a pair was masked during our masking procedure. A task counts as solved if all constraints in the sequence and the structure are satisfied, i.e. multiple candidates are valid for a given input task. We run each algorithm for one hour on each task of the benchmark in five independent runs and report the average number of solved tasks with the standard deviation around the mean. The results are shown in Figure 3. We observe that libLEARNA clearly outperforms antaRNA and MoiRNAiFold, indicating that our training strategy enabled libLEARNA to efficiently use the restrictions to find good regions in the design space.

**Figure 3.**
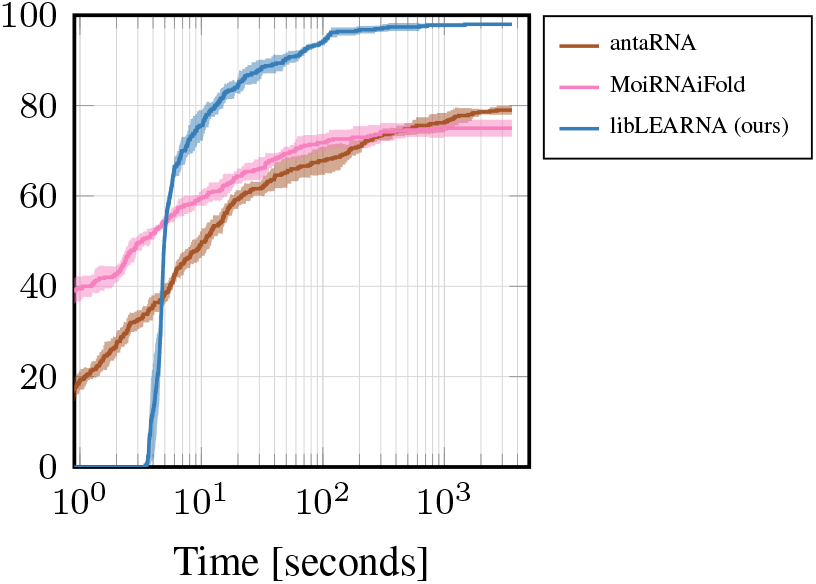
Comparison on randomly masked tasks with balanced brackets.

#### 4.1.3 Robustness to Changes of the Objective

The training objective of libLEARNA has three components: a folding algorithm for folding the candidate solution, a distance measure to quantify the difference between the folded candidate and the desired structure constraints (see Equation 12), and a reward function that uses the structure loss to define a reward signal (see Equation 14). To analyze the behavior of libLEARNA to changes in the objective, we change one component at a time.

##### libLEARNA Adapts to Changes of the Loss Function

To analyze the adaptation to changes of the loss function, we either change the distance measure, the folding algorithm, or both. libLEARNA is based on the Meta-LEARNA-Adapt approach [Runge et al., 2019] which uses a restart option that resets the weights of the policy network to the initial values every 1800 seconds. However, some changes might require a longer time for adaptation. We, therefore, evaluate each variant with and without the restart option. We again use the Eterna100 benchmark version 2 for evaluation. For the distance metric, we replace the Hamming distance with a Weisfeiler-Lehman graph kernel (WL) as recently proposed for the evaluation of secondary structure prediction [Runge et al., 2023]. This change affects Equation 13 by replacing the Hamming distance d_H_ with a WL distance measure *d*_*W L*_ = (1 − *S*_*W L*_), where *S*_*W L*_ is the similarity score obtained from the WL computations. For the folding engine, we use the RNAFold variant for the maximum expected accuracy (MEA) structure instead of the minimum free energy (MFE) structure, i.e., we replace the folding algorithm ℱ_MFE_ with ℱ_MEA_ in Equation 12. The accumulated number of solved tasks across all runs for each variant is shown in Table 2. Surprisingly, changing the distance measure results in better performance compared to the original libLEARNA baseline with and without restarts of the algorithm. However, for the folding engine, we observe that the change to the MEA folding objective decreases the performance in the setting with restarts, while it seems to be beneficial when there is no restart performed, independent of the distance measure. We conclude that changing the folding engine is a critical change to the objective that requires more time for adaptation. However, overall the performance of libLEARNA appears robust to changes in the loss function.

**Table 2:**
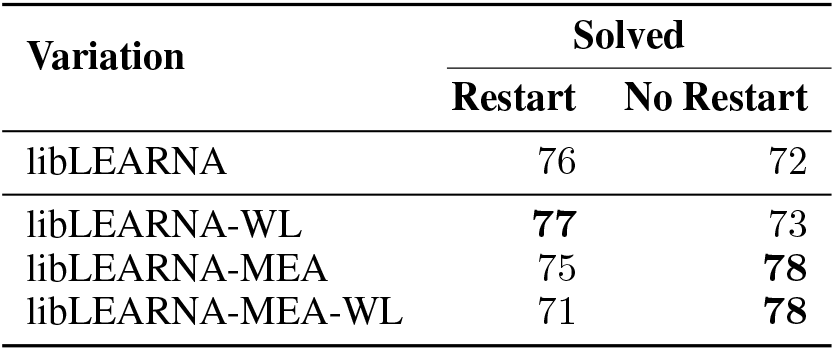
Analysis of changes applied to the loss function.

#### libLEARNA Adapts to Changes of the Reward Function

To analyze the influence of a change on the reward function, we evaluate libLEARNA when adding a GC-content objective. We implement an additional GC-loss term in the reward function. The GC-loss, *L*_*GC*_, is defined as the absolute deviation of the GC-content of the designed sequence *ϕ*, GC_*ϕ*_, from the desired GC-content, GC_desired_, allowing a given tolerance *ϵ*:

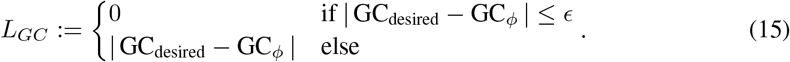

We use a tolerance *ϵ* = 0.01 for all experiments with desired GC-contents. For the reward function ℛ^*T*^ we then use the weighted sum of the structure-loss and the GC-loss:

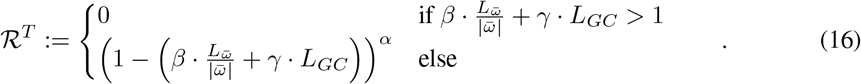

We set *β* = *γ* = 1 without further tuning and note that libLEARNA was not trained to include desired GC contents. Runge et al. [2019] propose a local improvement step (LIS) to aid the agent in solving a task once it is close to a solution. We adopt this procedure and additionally implement a GC-improvement step (GIS) that becomes active whenever the LIS is active. We describe the GIS in more detail in Appendix C. We evaluate libLEARNA against antaRNA [Kleinkauf et al., 2015] and MCTS-RNA [Yang et al., 2017] on all pseudoknot-free tasks of the ArchiveII dataset [Sloma and Mathews, 2016] with a timeout of one hour and report the accumulated number of solved tasks across five independent runs. Table 3 shows the results. Remarkably, libLEARNA is on par with MCTS-RNA while clearly outperforming antaRNA. This confirms our previous observation that libLEARNA can adapt to changes in the objective.

**Table 3:** Results on tasks with desired GC-contents of the ArchiveII dataset.

| Model | Solved |
| --- | --- |
| antaRNA | 52% |
| MCTS-RNA | **73%** |
| libLEARNa (ours) | **73%** |

### 4.2 Solving the Partial RNA Design Problem

In this section, we tackle the problem of RNA design for partially restricted design spaces of variable length, essentially solving the partial RNA design problem with varying objectives. We design theophylline riboswitch constructs using the training objective described in Equation 14 and the combined objective to design RNAs with desired GC-contents described in Equation 16, before we assess the performance of libLEARNA under completely new objectives.

#### 4.2.1 Automated Design of Theophylline Riboswitch Candidates

In this section, we generate diverse, variable-length theophylline riboswitch constructs in a single run of libLEARNA, following a previously published protocol [Wachsmuth et al., 2012]. Originally, Wachsmuth et al. [2012] constructed theophylline riboswitch candidates for transcriptional activation from (1) the TCT8-4 theophylline aptamer sequence and structure, (2) a spacer sequence of 6 to 20 nucleotides (nt), (3) a sequence of 10nt to 21nt complementary to the 3^*′*^-end of the aptamer, and (4) a U-stretch of 8nt at the 3^*′*^-end of the construct. To generate candidate sequences, Wachsmuth et al. [2012] create a large library of random sequences for the spacer region (6-20nt) and a library of sequences complementary to the 3^*′*^-end of the aptamer (10-21nt). From these sets, randomly sampled sequences were combined with the aptamer and the 8-U-stretch.

For the creation of the search space, we use the shared sequence and structure motifs of the six proposed riboswitch constructs by Wachsmuth et al. [2012] and combine them into a single design space formulation. The final definition of the design space and the proposed constructs of Wachsmuth et al. [2012] are shown in Table 7 in Appendix D.

**Table 4:** Overview of designed riboswitch candidates.

| Method | Valid Candidates [%] | Unique Structures |
| --- | --- | --- |
| Wachsmuth et al. [2012] | 41,7 | 6316.6 |
| *libLEARN*A | **70,9** | **8269.6** |

**Table 5:** Overview over the Datasets.

| Data Set | Tasks | Mean Length | Median Length | Min/Max Length |
| --- | --- | --- | --- | --- |
| Training1 (“long”) | 100000 | 470.9 | 300 | 200–8033 |
| Training2 (“short”) | 100000 | 103.9 | 99 | 23–200 |
| Training3 (“random”) | 100000 | 142.7 | 106 | 23–6361 |
| Validation | 100 | 135.8 | 94 | 46–1802 |

**Table 6:** Configuration space and final parameters of libLEARNA.

| Parameter Name | Type | Range | Prior | libLEARNNA |
| --- | --- | --- | --- | --- |
| state radius $\kappa$ | integer | [0, 32] | uniform | 10 |
| individual state composition $\sigma$ | categorical | [“target”, “design”] | uniform | “target” |
| action semantics | categorical | [“pair”, “single”] | uniform | “pair” |
| reward exponent $\alpha$ | float | [1, 12] | uniform | 10.76 |
| filter size in 1<sup>st</sup> conv layer | integer | $\{0\} \cup \{3, 5, \dots, 17\}$ | uniform | 0 |
| filter size in 2<sup>nd</sup> conv layer | integer | $\{0, 3, 5, 7, 9\}$ | uniform | 0 |
| # filter in 1<sup>st</sup> conv layer | integer | [1, 32] | log-uniform | 17 |
| # filter in 2<sup>nd</sup> conv layer | integer | [1, 32] | log-uniform | 24 |
| # LSTM layers | integer | [0, 3] | uniform | 0 |
| # units in every LSTM layer | integer | [1, 64] | log-uniform | 20 |
| # fully connected layers | integer | [1, 2] | uniform | 2 |
| # units in fully connected layers | integer | [8, 64] | log-uniform | 12 |
| embedding dimensionality | integer | [0, 21] | uniform | 17 |
| batch size | integer | [32, 256] | log-uniform | 247 |
| entropy regularization | float | $[1 \cdot 10^{-7}, 1 \cdot 10^{-2}]$ | log-uniform | $4.46 \cdot 10^{-7}$ |
| learning rate for PPO | float | $[1 \cdot 10^{-6}, 1 \cdot 10^{-3}]$ | log-uniform | $5.9 \cdot 10^{-4}$ |
| training data | categorical | [“random”, “short”, “long”] | uniform | “short” |
| training curriculum | categorical | [“random”, “sorted”] | uniform | “sorted” |

**Table 7:**
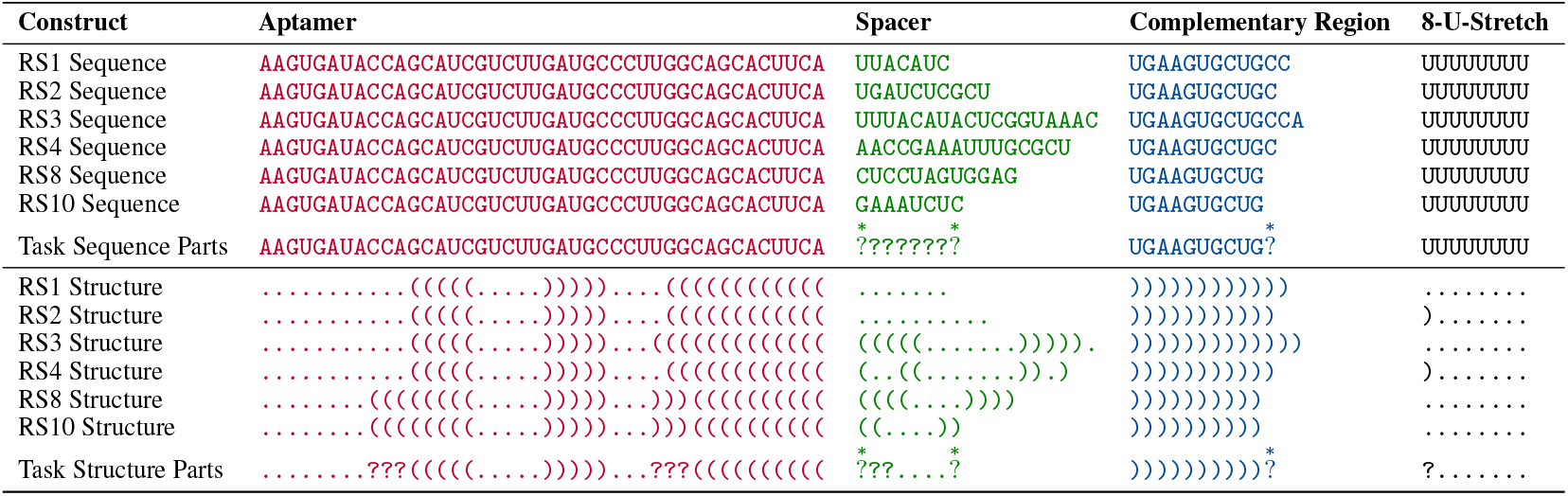
Originally proposed theophylline riboswitch constructs and partial RNA design space formulation. (Top) The sequence parts of the six originally proposed riboswitch constructs by Wachsmuth et al. [2012] and the sequence part of the design space. (Bottom) The corresponding structure parts of the constructs and the structural part of the desiDesign Space of libLEARNA for the design of theophylline riboswitch constructs. The constructs RS1, RS2, RS3, RS4, RS8 and RS10 were proposed by (37). Highlighted regions mark parts that are shared across all the original riboswitch constructs. These regions are used to construct a design space for libLEARNA. At the bottom, the figure shows an example prediction of libLEARNA for the given design space.gn space. Red: TCT8-4 theophylline aptamer; green: variable length spacer domain; blue: domain ought to pair with the aptamer (complementary to the 3^*′*^-end of the aptamer sequence in the original design); black: 8-U-stretch. Masked positions are indicated with `?`, positions for extensions are indicated with 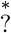.

We assess the performance of libLEARNA against the original library generation procedure proposed by Wachsmuth et al. [2012] in two experiments: (1) The design of a riboswitch library based on sequence and structure constraints only, and (2) the design of sequences when additionally querying libLEARNA to design candidates with a specific GC-content, given a tolerance of 0.01. For each experiment, we generate 50000 candidates with the approach of Wachsmuth et al. [2012] and libLEARNA with five random seeds and evaluate them using the same protocol as used by Wachsmuth et al. [2012] without ranking the constructs via z-scores.

We observe that libLEARNA generates considerably more candidates that pass the design criteria compared to the original procedure proposed by Wachsmuth et al. [2012], yielding 71% satisfying candidates on average, compared to 42% for the original procedure (Table 4). Further, the candidates are nearly uniformly distributed across the lengths of the design space, especially for the longer sequences (Figure 4 left), and the structure diversity generated by libLEARNA is around 23% higher (Table 4). When designing candidates with desired GC-contents, libLEARNA provides up to 47% more candidates that satisfy the design criteria and the desired GC-content (Figure 4). Remarkably, libLEARNA can also design candidates on the margins of possible GC-contents (0.3 and 0.6) which is between 0.29 and 0.63 for the given riboswitch design space.

**Figure 4.**
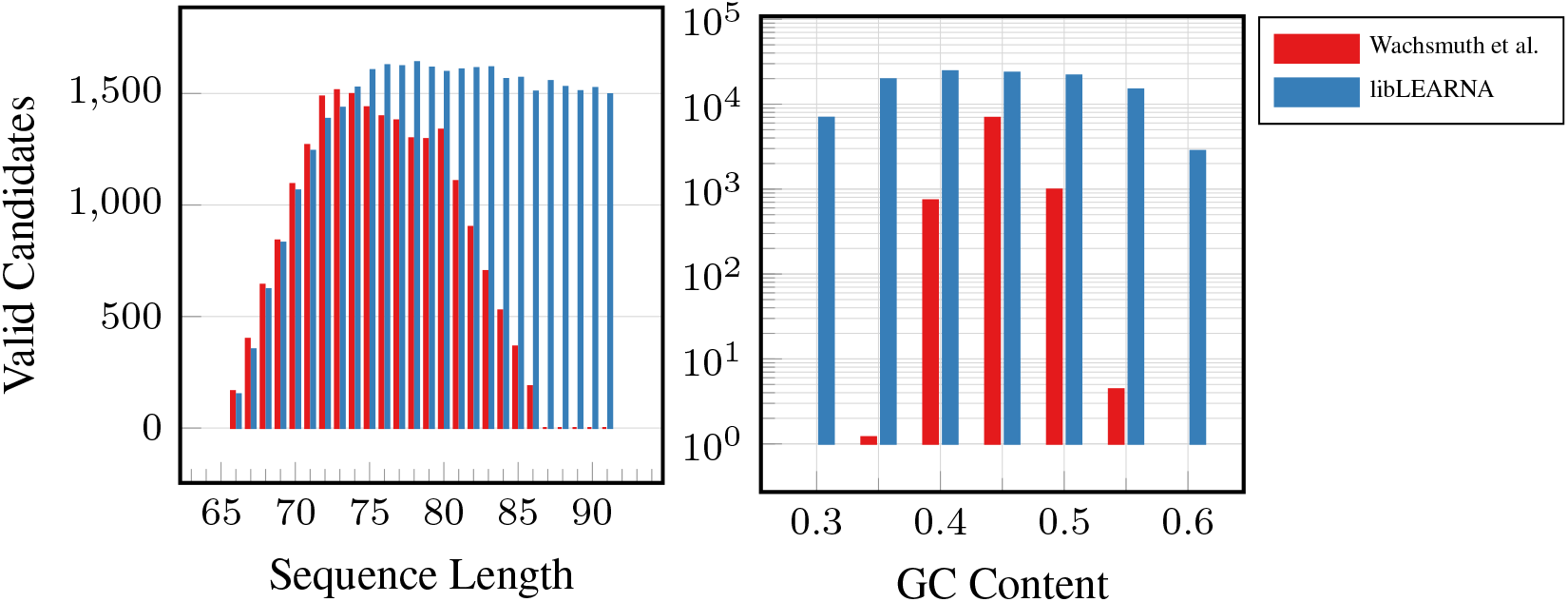
Average number of designed riboswitches across different lengths and GC-contents.

#### 4.2.2 libLEARNA Adapts to New Objectives

Partial RNA design allows the definition of arbitrary objectives. We, therefore, assess libLEARNA’s performance on two objectives that are independent of a folding algorithm. In the first experiment, we seek to design RNAs that belong to the family of Hammerhead ribozymes (Type III) (Rfam family: RF00008). To guide the search, we use the consensus structure and the conserved nucleotides provided by the Rfam database, while allowing for exploration at the beginning, end, and in the loop regions of the structure. We use the covariance model of the family (CLEN 54) to receive bitscores for each design, *ϕ*, using Infernal [Nawrocki and Eddy, 2013] and design candidates with a total length between 50 and 60 nucleotides. We implement a new reward function that directly optimizes for bitscores: *R*^*T*^ = *bitscore*(*ϕ*). The reward for a candidate that does not match the covariance model at all is set to − 200. We compare libLEARNA with a random agent that uniformly samples actions at each unconstrained position of the search space. The results are shown in Figure 6 in Appendix E. libLEARNA quickly optimizes the bitscore to roughly 30 on average, while the average bitscores of the random agent remains below zero. For reference, we sample 1000 sequences directly from the covariance model using cmemit of the Infernal package. The highest achieved bitscore of these sequences is 71.1 compared to 72.2 when designing sequences with libLEARNA. However, the search space definition might affect the performance. We, therefore, analyze the influence of the search space definition in a second experiment where we use libLEARNA to design candidates for RNA-RNA interactions (RRI). More precisely, we design candidate sequences for a given target RNA that minimize the energy of the interaction obtained from IntaRNA [Mann et al., 2017]. We use two RNAs of *Prochlorococcus marinus*: an mRNA target (GenBank Accession: BX548174; Region: 1069333..1069507) and an ncRNA (GenBank Accession: NC 008817; Region: 1002095..1002151) as template for the search. We define (1) A completely unrestricted search space that only contains a variable length wildcard symbol such that libLEARNA designs sequences guided solely by the reward function. (2) A search space that only restricts the structure part; We fold the example sequence and introduce a point of extension (a variable length wildcard symbol 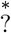 in the middle of the resulting two hairpins. (3) A search space with restrictions in the structure and the sequence part, by providing 10 nucleotides from the original sequence at random positions in addition to the search space defined in (2). We implement a new reward function that optimizes for the inverse of the energy: *R*^*T*^ = (*Energy* · (−1))^3^. Again we compare libLEARNA to its random variant in five runs for 500000 episodes. Figure 5 shows the results of the RRI design task while the energy is averaged across 500 steps and all runs. We observe that libLEARNA can design candidates with a lower energy of the complex compared to the random agent and the initial example sequence (indicated by a red line in Figure 5). Interestingly, libLEARNA quickly adapts to the new reward function even without any sequence and structure information (unconstrained). However, the structure information seems to be beneficial in the long run since the final energy for the designed candidates using the design space with the structure information only is the lowest of all three search space formulations. The search space that contains sequence and structure restrictions seems to impose more challenges to the design process. While the resulting candidates still clearly show a lower energy with the target RNA compared to the original sequence, the optimization is slower and results in a higher final energy. Overall, we conclude that the formulation of the search space can have a substantial impact on the outcome of a given design task.

**Figure 5.**
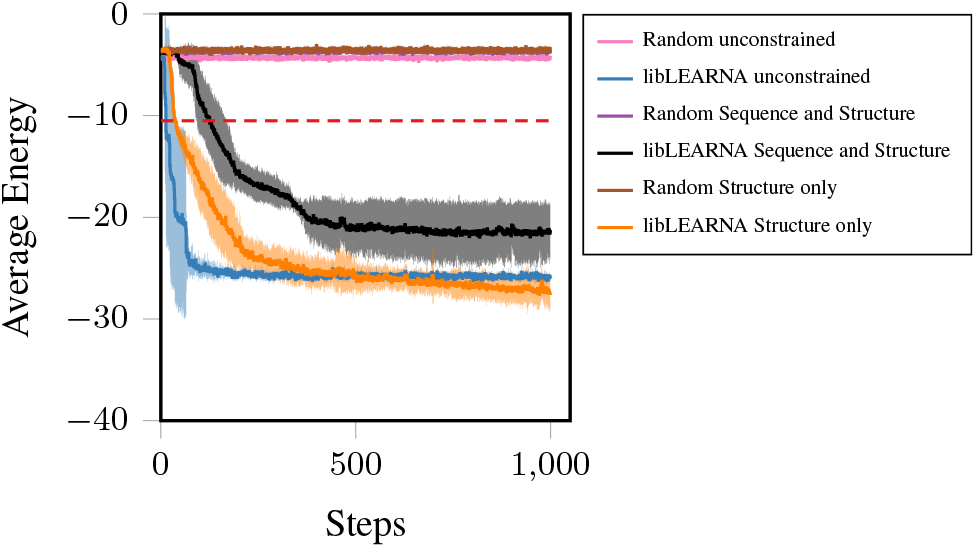
The average energy of RRI complex. A step in the figure corresponds to 500 episodes. The values are averages across five independent runs with standard deviation around the mean displayed as confidence bounds. The dashed red line indicates the energy level of the template sequence in complex with the target RNA.

**Figure 6.**
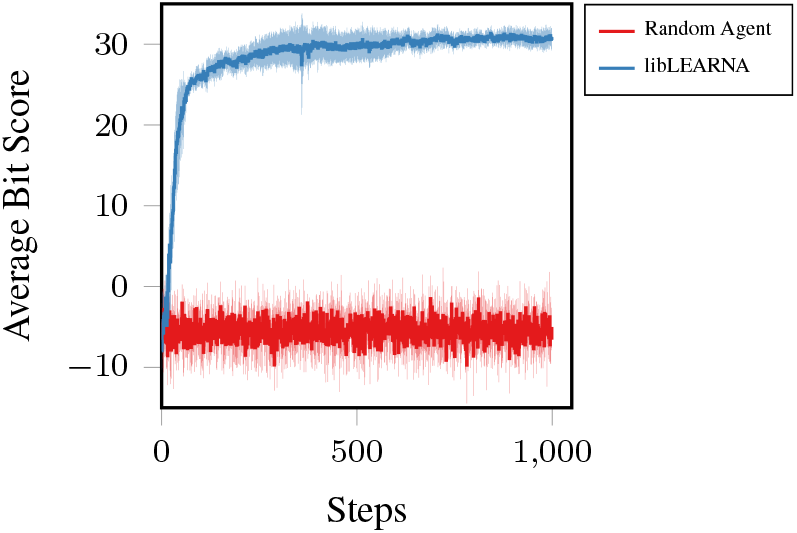
Average bitscore of designed candidates for the covarinance model of Hammerhead Ribozyme (Type III). A step in the figure corresponds to 100 episodes.

## 5 Conclusion

In this work, we propose *partial RNA design*, a new RNA design paradigm formulated as a constraint satisfaction problem. The idea is to treat RNA design as a search in a partially restricted design space that can be explored by a machine under different objectives. We then propose libLEARNA, a robust automated reinforcement learning algorithm, capable of navigating partially restricted design spaces. We show that libLEARNA has strong performance across a wide range of different RNA design tasks, including restrictive design settings with fixed lengths and tasks that require exploration of open design spaces and the generation of variable-length candidates. Our work describes a novel approach to RNA design where the algorithm provides large amounts of solutions, given the restrictions of its search space rather than designing one or few solutions for a very narrow task definition. We believe that our new approach adds a new dimension to the RNA design problem and that libLEARNA is a useful tool for future RNA design endeavors.

## Acknowledgments

We acknowledge funding by the Deutsche Forschungsgemeinschaft (DFG, German Research Foundation) under SFB 1597 (SmallData), grant number 499552394, and through grant number 417962828. We also acknowledge support by the state of Baden-Württemberg through bwHPC and the German Research Foundation (DFG) through grant no INST 39/963-1 FUGG. Finally, we acknowledge funding by the European Union (via ERC Consolidator Grant DeepLearning 2.0, grant no. 101045765). Views and opinions expressed are however those of the author(s) only and do not necessarily reflect those of the European Union or the European Research Council. Neither the European Union nor the granting authority can be held responsible for them.

## A Data

## B Optimization

## C Local Improvement Steps

Runge et al. [2019] propose a local improvement step (LIS) that exhaustively evaluates neighboring sequences once the agent is close to a solution. We kept the LIS cut-off parameter *ξ* unchanged as proposed at Runge et al. [2019], setting it to 5 which corresponds to at most 4^4^ = 256 neighboring sequences that are explored during LIS. For our experiments of RNA design with desired GC-contents we develop a similar GC-improvement procedure (GIS). The GIS becomes active whenever the LIS is active and is applied after the LIS. For the GIS we iteratively select a random nucleotide without replacement, while the respective choice depends on the current GC-content of the candidate sequence and the desired GC-content (either A or U if the GC content is too low or G and C if the GC-content is too high). We then replace the nucleotide with a randomly chosen nucleotide to either increase or decrease the GC-content. At each iteration, we fold the sequence and evaluate the structure-loss to keep it at least as low as at the beginning of the GIS. This procedure is repeated until no nucleotides are left for replacement. We provide pseudocode of the GIS in Algorithm 1.

### Algorithm 1

GC Improvement Step (GIS).

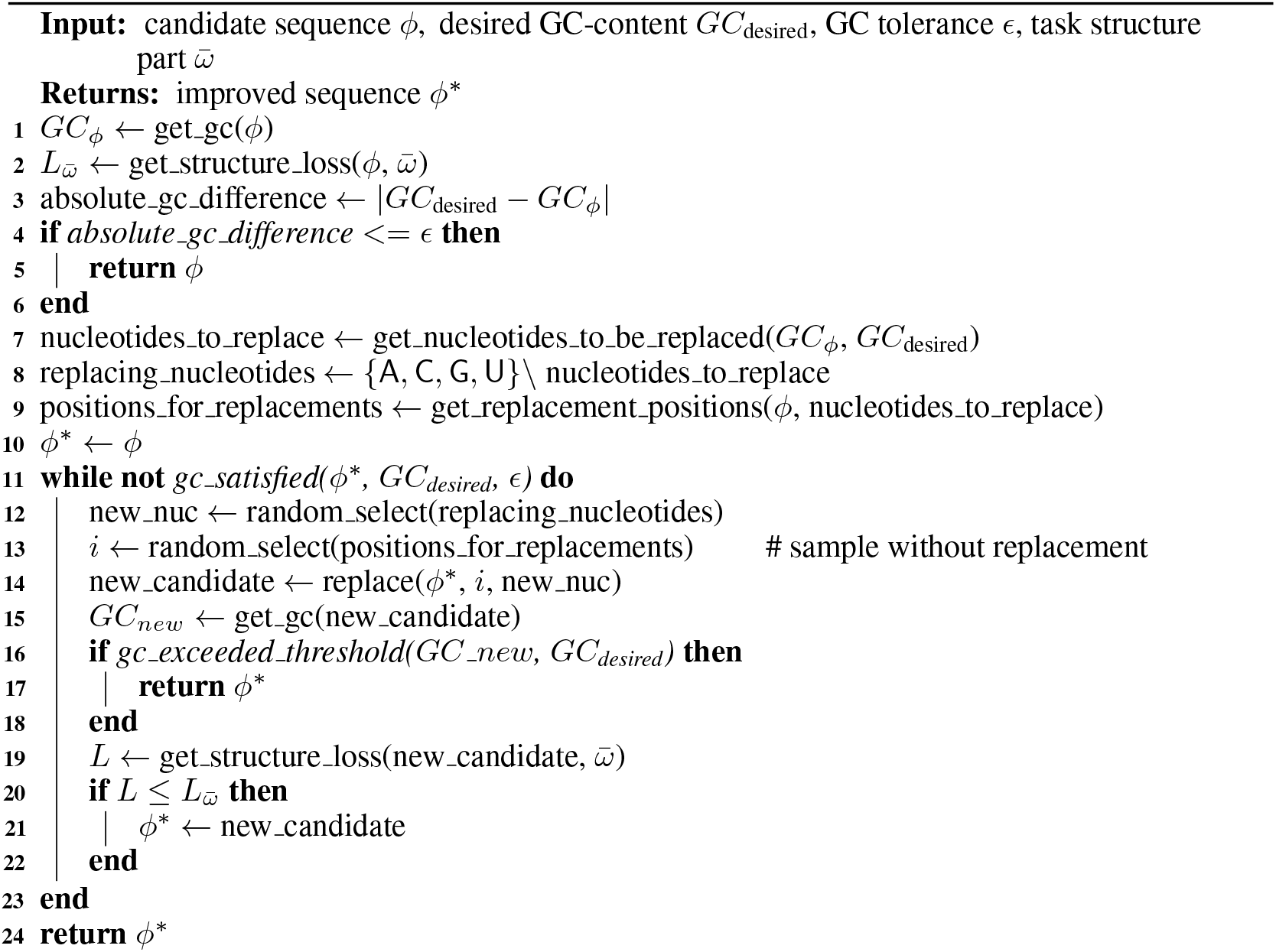

## D Riboswitch Design Space

## E Additional Results

## Notes

### Competing Interest Statement

The authors have declared no competing interest.

### Summary of Updates

Manuscript updated to the current version; under review.

## References

Stefan Hammer, Christian Günzel, Mario Mörl, and Sven Findeiß. Evolving methods for rational de novo design of functional rna molecules. Methods, 161:54 – 63, 2019. ISSN 1046-2023. Development and engineering of artificial RNAs.

Manja Wachsmuth, Sven Findeiß, Nadine Weissheimer, Peter F. Stadler, and Mario Mörl. De novo design of a synthetic riboswitch that regulates transcription termination. Nucleic Acids Research, 41(4):2541–2551, 12 2012. ISSN 0305-1048.

Mo Li, Mengxi Zheng, Siyu Wu, Cheng Tian, D. Liu, Yossi Weizmann, Wen Jiang, Guansong Wang, and Chengde Mao. In vivo production of rna nanostructures via programmed folding of single-stranded rnas. Nature communications, 9(1):1–9, 2018.

Yuri Nozawa, Megumi Hagihara, Md Sohanur Rahman, Shigeyoshi Matsumura, and Yoshiya Ikawa. Rational design of an orthogonal pair of bimolecular rnase p ribozymes through heterologous assembly of their modular domains. Biology, 8(3):65, 2019.

Francis E Reyes, Andrew D Garst, and Robert T Batey. Strategies in rna xdcrystallography. Methods in enzymology, 469:119–139, 2009.

Ivo Hofacker, Walter Fontana, Peter Stadler, Sebastian Bonhoeffer, Manfred Tacker, and Peter Schuster. Fast Folding and Comparison of RNA Secondary Structures. Monatshefte fuer Chemie/Chemical Monthly, 125:167–188, 02 1994.

Robert Kleinkauf, Torsten Houwaart, Rolf Backofen, and Martin Mann. antaRNA–Multi-objective inverse folding of pseudoknot RNA using ant-colony optimization. BMC bioinformatics, 16(1):389, 2015.

Frederic Runge, Danny Stoll, Stefan Falkner, and Frank Hutter. Learning to design RNA. In International Conference on Learning Representations, 2019.

Gerard Minuesa, Cristina Alsina, Juan Antonio Garcia-Martin, Juan Carlos Oliveros, and Ivan Dotu. Moirnaifold: a novel tool for complex in silico rna design. Nucleic acids research, 49(9):4934– 4943, 2021.

Mirela Andronescu, Anthony P Fejes, Frank Hutter, Holger H Hoos, and Anne Condon. A new algorithm for rna secondary structure design. Journal of molecular biology, 336(3):607–624, 2004.

Jessica S Reuter and David H Mathews. Rnastructure: software for rna secondary structure prediction and analysis. BMC bioinformatics, 11(1):1–9, 2010.

Assaf Avihoo, Alexander Churkin, and Danny Barash. Rnaexinv: An extended inverse rna folding from shape and physical attributes to sequences. BMC bioinformatics, 12(1):319, 2011.

Matan Drory Retwitzer, Vladimir Reinharz, Yann Ponty, Jérôme Waldispühl, and Danny Barash. incaRNAfbinv: a web server for the fragment-based design of RNA sequences. Nucleic Acids Research, 44(W1):W308–W314, 05 2016. ISSN 0305-1048.

Juan Antonio Garcia-Martin, Peter Clote, and Ivan Dotu. Rnaifold: a constraint programming algorithm for rna inverse folding and molecular design. Journal of Bioinformatics and Computational Biology, 11(02):1350001, 2013.

Juan Antonio Garcia-Martin, Ivan Dotu, and Peter Clote. Rnaifold 2.0: a web server and software to design custom and rfam-based rna molecules. Nucleic Acids Research, 43(W1):W513–W521, 2015.

Peter Fletcher, Hughes Hoyle, and C. Wayne Patty. Foundations of discrete mathematics. Boston, MA: PWS-Kent Publishing Company, 1991. ISBN 0-534-92373-9.

Jacob Devlin, Ming-Wei Chang, Kenton Lee, and Kristina Toutanova. Bert: Pre-training of deep bidirectional transformers for language understanding. arXiv preprint arXiv:1810.04805, 2018.

Jack Parker-Holder, Raghu Rajan, Xingyou Song, André Biedenkapp, Yingjie Miao, Theresa Eimer, Baohe Zhang, Vu Nguyen, Roberto Calandra, Aleksandra Faust, Frank Hutter, and Marius Lindauer. Automated reinforcement learning (autorl): A survey and open problems. Journal of Artificial Intelligence Research (JAIR), 74:517–568, 2022.

Stefan Falkner, Aaron Klein, and Frank Hutter. BOHB: Robust and efficient hyperparameter optimization at scale. In Jennifer Dy and Andreas Krause, editors, Proceedings of the 35th International Conference on Machine Learning, volume 80 of Proceedings of Machine Learning Research, pages 1437–1446, Stockholmsmässan, Stockholm Sweden, 10–15 Jul 2018. PMLR.

Ronny Lorenz, Stephan H. Bernhart, Christian Hönerzu Siederdissen, Hakim Tafer, Christoph Flamm, Peter F. Stadler, and Ivo L. Hofacker. Viennarna package 2.0. Algorithms for Molecular Biology, 6(1):26, Nov 2011. ISSN 1748-7188.

Sam Griffiths-Jones, Alex Bateman, Mhairi Marshall, Ajay Khanna, and Sean R. Eddy. Rfam: an RNA family database. Nucleic Acids Research, 31(1):439–441, 01 2003. ISSN 0305-1048. doi: 10.1093/nar/gkg006.

Peter Henderson, Riashat Islam, Philip Bachman, Joelle Pineau, Doina Precup, and David Meger. Deep reinforcement learning that matters. In Thirty-Second AAAI Conference on Artificial Intelligence, 2018.

Rohan V Koodli, Boris Rudolfs, Hannah K Wayment-Steele, Eterna Structure Designers, and Rhiju Das. Redesigning the eterna100 for the vienna 2 folding engine. bioRxiv, pages 2021–08, 2021.

Frederic Runge, Jörg K. H. Franke, Daniel Fertmann, and Frank Hutter. Rethinking performance measures of rna secondary structure problems, 2023.

Xiufeng Yang, Kazuki Yoshizoe, Akito Taneda, and Koji Tsuda. Rna inverse folding using monte carlo tree search. BMC bioinformatics, 18(1):468, 2017.

Michael F Sloma and David H Mathews. Exact calculation of loop formation probability identifies folding motifs in rna secondary structures. RNA, 22(12):1808–1818, 2016.

Eric P Nawrocki and Sean R Eddy. Infernal 1.1: 100-fold faster rna homology searches. Bioinformatics, 29(22):2933–2935, 2013.

Martin Mann, Patrick R Wright, and Rolf Backofen. Intarna 2.0: enhanced and customizable prediction of rna–rna interactions. Nucleic acids research, 45(W1):W435–W439, 2017.

